# Free energy and stacking of eumelanin nanoaggregates

**DOI:** 10.1101/2021.08.31.458381

**Authors:** Sepideh Soltani, Shahin Sowlati-Hashjin, Conrard Giresse Tetsassi Feugmo, Mikko Karttunen

## Abstract

Eumelanin, a member of the melanin family, is a black-brown insoluble pigment. It possesses a broad range of properties such as antioxidation, free radical scavenging, photo-protection, and charge carrier transportation. Surprisingly, the exact molecular structure of eumelanin remains undefined. It is, however, generally considered to consist of two main building blocks, 5,6-dihydroxyindole (DHI) and 5,6-dihydroxyindole carboxylic acid (DHICA). We focus on DHI and report, for the first time, a computational investigation of structural properties of DHI eumelanin aggregates in aqueous solutions. First, multi-microsecond molecular dynamics (MD) simulations at different concentrations were performed to investigate aggregation and ordering of tetrameric DHI-eumelanin protomolecules. This was followed by umbrella sampling (US) and density functional theory (DFT) calculations to study the physical mechanisms of stacking. Aggregation occurs through formation of nanoscale stacks and was observed in all systems. Further analyses showed that aggregation and coarsening of the domains is due to decrease in hydrogen bonds between the eumelanins and water; while domains exist, there is no long-range order. The results show non-covalent stacks with and interlayer distance between eumelanin protomolecules being less than 3.5 Å. This is in good agreement with transmission electron microscopy data. Both free energy calculations and DFT revealed strong stacking interactions. The electrostatic potential map provides an explanation and a rationale for the slightly sheared relative orientations and, consequently, for the curved shapes of the nanoscale domains.

## INTRODUCTION

Melanins are a family of dark insoluble biological pigments found in many organisms such as fungi and bacteria and in different plant, animal and human tissues^1–3^. In human bodies, melanins are present in eye, skin, hair, brain, and inner ear (cochlea). In the eye, melanin-containing cells are localized in the uveal tract (iris, ciliary body), choroid, and in the retinal pigment epithelium. Melanins have a very broad range of desirable properties including antioxidant^4,5^, free radical scavenging^6^, photo-protection^7–10^ and charge carrier transport^11,12^, and they have also been associated with diseases such as Parkinson’s^13–15^, age-related macular degeneration^16–18^, age-related/noise-induced hearing loss ^19^ and skin diseases such as albinism, vitiligo and malignant melanoma^8,20,21^. These properties have lead to a growing interest in application fields such as bioelectronics, batteries^22,23^, biomimetic interfaces^24,25^, ophthalmic drug development^26–28^ and bioimaging^10,23^.

Melanin is usually classified into three groups, eumelanin, pheomelanin and neuromelanin, but sometimes allomelanin and pyomelanin are also listed as separate classes^29^. Our focus is on eumelanin which has two naturally occurring major forms, 5,6-dihydroxyindole-2-carboxylic acid (DHICA) and 5,6-dihydroxyindole (DHI), see Fig. 1a. Natural eumelanin is a brown-black insoluble photoprotective pigment of human skin and eyes^30,31^. Its characteristic features originate from the high number of carboxylic acid residues with negative charge^32^, the number of aromatic rings with *π*-interactions^33^, hydrogen bonding and van der Waals interactions^32,34^. The proportions of the two components (DHICA and DHI) have significant variations depending on the source where melanin was extracted. It has been reported that in rodent hair and melanoma, the DHICA form may have proportions in the range of 59-98%^35^ and in humans the amount of DHICA has been reported to be in the range of 19-55% depending on tissue and sample.^35–37^ In applications, typically synthetic eumelanin containing mostly DHI is used.^31,38^

**Figure (1).**
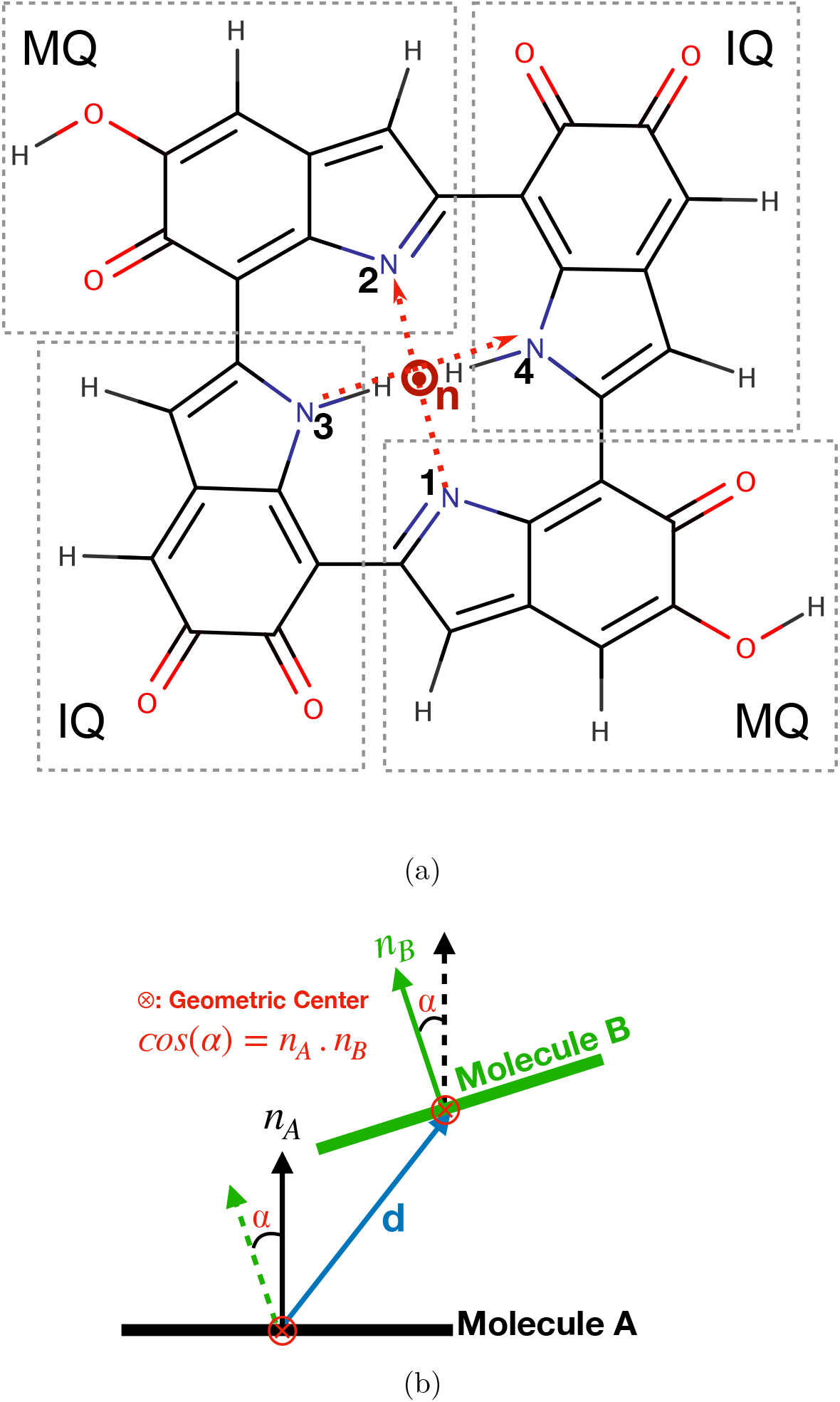
a) Chemical structure of tetrameric DHI-eumelanin based on the model of Kaxiras et al.^52^. Building blocks of tetrameric DHI-eumelanins are indolequinone (IQ) and its quinone-methide tautomer (MQ). Definition of the normal vector *n* (pointing out of the plane) is also shown. b) Geometric center distance *d* between two tetrameric DHI-eumelanin protomolecules. *n*_*A*_ and *n*_*B*_ are normal vectors corresponding to molecules A and B. Stacks are parallel, if *α* < 18.2° and *d* < 7.0 Å.

The molecular structure of eumelanin has been a challenge, and D’Ischia et al.^39^ have pointed out that there is no single well-defined structure for eumelanin. They underline this by stating that “*…it is not possible to provide an accurate picture beyond a statistical description of main units and functional groups.”* For larger aggregates, experiments have shown that sheets of protomolecules may stack to form onion-like nanostructures with an interlayer distance of between 3.3 and 4.0 Å^40,41^. Computer simulations have also shown non-covalent *π* – *π* stacking in heteroaromatic systems^42–47^. Majority of computational investigations are on the porphyrin-like planar tetramer DHI-eumelanin ^42,43,48–51^.

The basis for most of computer simulations of DHI eumelanin is the model of Kaxiras et al.^52^ who used DFT calculations to develop a model for tetrameric DHI-eumelanin. The model was constructed such that it reproduces the following key properties: 1) The small size (15 – 20 Å) of molecules corresponding to tetramers or pentamers, 2) the molecules must have the ability to be stacked in a planar graphite-like arrangement, 3) the ability of capturing and releasing one metal ion without morphological changes, and 4) broad absorption in UV-visible spectrum.

In this study, we used the tetrameric DHI-eumelanin structure of Kaxiras et al. There are several unresolved questions regarding how eumelanin aggregates form individual protomolecules in aqueous solutions and how the aqueous environment affects eumelanin protomolecules. Currently, there are only a few MD studies that have focused on monomers, nanocrystals (no water present) or near dry systems, and typical simulation times are less than 5 ns^42,43,46,47,51^. Here, our motivation is to investigate the effect of an aqueous environment on eumelanin molecules and their behavior.

## METHODOLOGY

### Tetrameric DHI-eumelanin model

The tetrameric DHI-eumelanin model (Fig. 1a) was originally proposed by Kaxiras et al. based on a molecular structure consisting of four monomer units with an inner porphyrin ring^52^. Later, Meng and Kaxiras^53^ used time-dependent density functional theory (DFT) calculations and found (in the absence of water) that two tetrameric DHI-eumelanins with *π*-stacking and van der Waals interactions can arrange to a quasi-parallel structure with interlayer distance of 3.0 – 3.3 Å. They also suggested that two tetrameric DHI-eumelanins can bind in a covalent manner through interlayer C-C bonds.

Chen et al.^42,43^ used the model of Kaxiras et al. ^52,53^ to investigate what they called ‘dry’ and ‘wet’ systems. ‘Dry’ systems had no water and ‘wet’ systems had 16 water molecules per tetrameric DHI-eumelanin. They observed stacked planes with interlayer distance around 3.5 – 4.0 Å (minimum distance) and since the planes were slightly tilted, the distance between the two centers of geometry was usually in the range of about 6.0 Å. Henceforth, we use eumelanin term instead of tetrameric DHI-eumelanin protomolecule and DHI notation in figures for simplicity.

### Geometry Optimization and Parameterization

In the current study, geometry optimization was done using the Gaussian 09^54^ software following the procedure of Kaxiras et al.^52^ The Becke three-parameter Lee-Yang-Parr (B3LYP)^55–60^ functional was adopted together with the split valence (polarized) basis set 6-311G(d) ^61–63^. The Pople basis sets such as 6-311G(d) are usually used for organic molecules^63^. The partial atomic charges were obtained from the Gaussian 09 software with the electrostatic potential (ESP)^64,65^ fit at the same level of theory. Other parameters for bonded and non-bonded interactions were collected using LigParGen server^66^ which is compatible with the Optimized Potentials for Liquid Simulations for All Atoms (OPLS-AA)^67,68^ force field. The parameters for eumelanin are shown in Table S1.

Dimeric eumelanins systems were fully optimized at IEF - PCM - CAM - B3LYP / 6-311G (d,p) level of theory (*ϵ* = 78.36; water) as implemented in Gaussian 16 suite of programs and the corresponding energies were evaluated with three different functional/ basis set combinations, namely, IEF - PCM - M06 - 2X / 6-311++G (2df,2pd), IEF - PCM - CAM - B3LYP - D3BJ / 6-311 ++ G(2df,2pd), and IEF - PCM - B3LYP - D3BJ / 6 - 311 ++G (2df,2pd).

The M06-2X functional^69^ is often recommended for non-covalent interactions and CAM-B3LYP^70^ includes long-range corrections. In B3LYP-D3BJ, the D3 version of Grimme’s dispersion has been added with Becke-Johnson damping^71^. The dimer with the highest energy (i.e., twisted) was selected as the reference energy and the energies of other dimeric conformations are reported relative to the reference.

### Molecular Dynamics Simulations

MD simulations of eumelanin in aqueous solutions with varying concentrations were performed using the Gromacs 2019.3 package.^72^ For water, the SPC/E model (extended simple point charge)^73^ was used and periodic boundary conditions were applied in all directions. The Lennard-Jones and the real space electrostatic interactions were cut off at 1.0 nm, and the long-range electrostatic interactions were handled by the particle-mesh Ewald (PME) method^74,75^ with cubic interpolation of order four.

Eumelanins and water were separately coupled to the heat bath at temperature of 300 K using the V-rescale algorithm^76^ with coupling constant of 0.1 ps. Initial equilibration was performed in the NVT (constant particle number, volume and temperature) ensemble for 2 ns. This was followed by pre-equilibration simulations in the NPT (constant particle number, pressure, and temperature) ensemble using the Parrinello-Rahman barostat^77^ at 1 bar with compressibility of 4.5 × 10^−5^ bar^−1^ and time constant of 2.0 ps for 2.0 ns. For production runs the bond lengths were kept constant with the P-LINCS algorithm^78^. All systems were charge neutral and the number of eumelanin molecules ranged from two to 343. Except for the two largest systems, two replicas of each system were simulated. All systems and simulations times are listed in Table 1. Visualizations were done using Visual Molecular Dynamics (VMD)^79^ and PyMol.^80^

**Table (1).** Eumelanin systems: Different initial conformations

| Number of eumelanins | Number of water molecules | Box Size (nm<sup>3</sup>) | Time ns |
| --- | --- | --- | --- |
| 2 neutral | 3,950 | (4.93)<sup>3</sup> | 1,207 |
| 2 neutral | 8,925 | (6.50)<sup>3</sup> | 1,000 |
| 4 neutral | 3,975 | (4.95)<sup>3</sup> | 2,200 |
| 4 neutral | 3,960 | (4.95)<sup>3</sup> | 2,100 |
| 8 neutral | 3,843 | (4.93)<sup>3</sup> | 2,070 |
| 8 neutral | 4,307 | (5.11)<sup>3</sup> | 2,620 |
| 27 neutral | 12,973 | (7.40)<sup>3</sup> | 2,080 |
| 27 neutral | 14,789 | (7.72)<sup>3</sup> | 2,000 |
| 64 neutral | 44,537 | (11.66)<sup>2</sup> × 10.14 | 1,300 |
| 343 neutral | 65,541 | (17.30)<sup>3</sup> | 244 |

### Free-Energy Calculations

Umbrella sampling (US)^81^ with the Weighted Histogram Analysis Method^82^ (WHAM) were used to calculate the potential of mean force (PMF)^83^ for binding interactions between two parallel stacked eumelanins. The initial structures were taken from the coordinates of the MD simulation of eight eumelanins at 2 *μs* with four different face orientations as shown in Fig. 2. The pathway for each system was divided into 23 windows with the reaction coordinate values (distance between two center of mass (COM)) separated by 0.3 Å.

**Figure (2).**
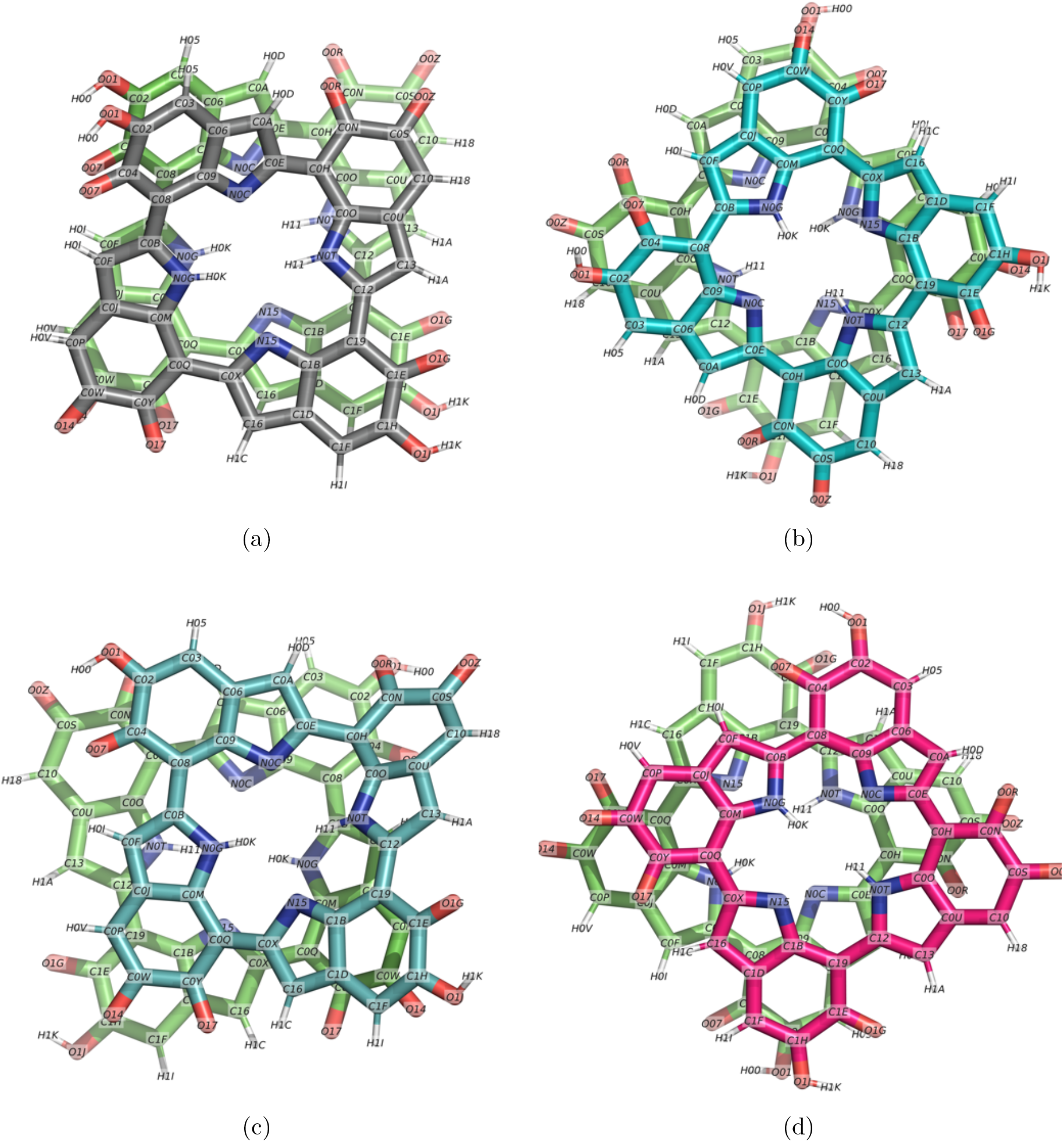
Snapshots of two eumelanins parallel staggered geometry in different rotational orientations. a) exact same order b) same order and 90° rotation c) mirror/flip d) mirror/flip and 90° rotation.

Pulling rate of 0.001 nm/ps was applied. The harmonic force constant was set to 1000 kJ/(mol × nm^2^) and 20 ns was used for each window to compute the PMF profile. Bias force was applied to the COM of an eumelanin in the *y*-direction (perpendicular to the eumelanin plane) and the other eumelanin molecule was maintained at a fixed location. The systems were subjected to a series of 24 simulation windows with 0.3 Å increments to provide the PMF profile as a function of the distance.

## RESULTS AND DISCUSSION

### Aggregation behavior

We start the discussion of results by focusing on aggregation behavior. We divide the systems into two categories, small and intermediate to large systems. The small systems show the basic behavior, and the intermediate ones provide the crossover to the complex aggregation seen in large systems. As the first quantity, we discuss the angle distribution, Fig. 3. The figures show two curves: One (black) for the average over the first 100 ns of the simulation and the other (red) for the last 100 ns of the simulation. Since the systems were started with random initial conditions, this way of plotting shows the evolution of the distribution toward equilibrium.

**Figure (3).**
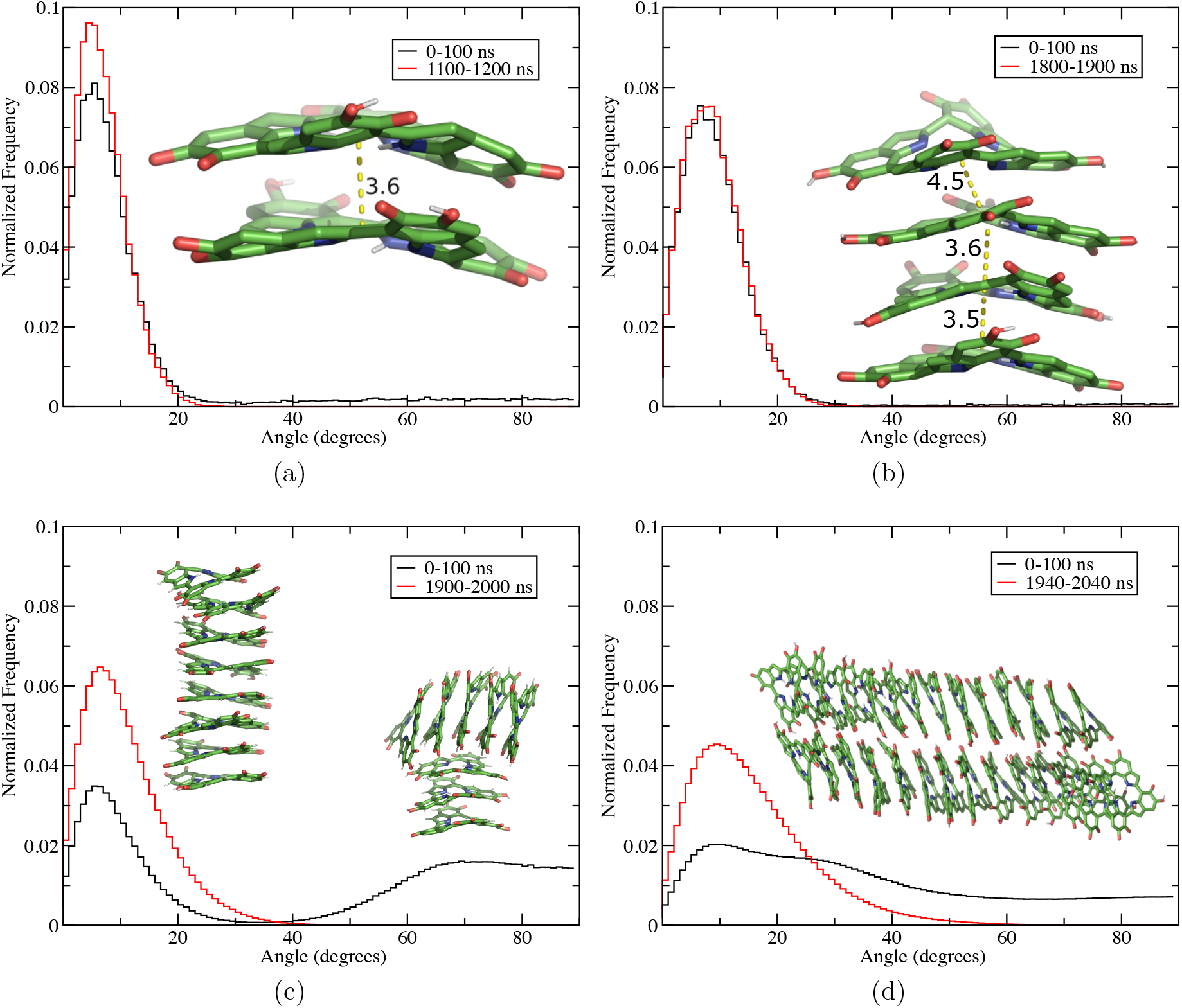
Comparison of angular distribution between eumelanins during the first 100 ns from a random initial conformation and the last 100 ns in aqueous bulk solution. a) two eumelanins, b) four eumelanins, c) eight eumelanins, d) 27 eumelanins. The yellow lines show the distance between the COGs of eumelanins, see Fig. 1 for the definition.

### Small systems

The smallest system has two eumelanin molecules, see Table 1 for system details. When two eumelanins were placed in water environment (random position and orientation), they aggregated and stacked in ≈ 11 ns. The timescale was the same in independent repeated simulations. As Fig. 3a shows, the angle distribution (see Fig. 1 for definitions) was established already within the first 100 ns. The inset shows the stacked configurations the molecules take, the center of geometry (COG) eumelanin-eumelanin distance and how the protomolecules bend in water environment.

The formation of stacks is consistent with Chen et al.^42,43^. We use their definition to evaluate the distance between the layers and to find when and how eumelanin stacks are formed during the simulations. Two eumelanins belong to a same secondary structure, namely stacked, if the following two criteria are met: 1) The distance *d* between their COGs is smaller than 7.0 Å. To find the COG of a eumelanin molecule, we average over the coordinates of the four nitrogen atoms in the interior ring as shown in Fig. 1a. 2) The angle between the normal vectors of two eumelanins must be smaller than 18.2°, see Fig. 1b for the definitions. Increasing the system size to four eumelanins did not lead to any significant changes, Fig. 3b. The molecules stacked again very rapidly in ≈ 10 ns as in the case of two eumelanins. The distribution becomes slightly broader and the peak has a small but noticeable shift toward larger values.

Figure 3c shows the evolution of the angle distribution for eight eumelanins. Similar to the previous cases, stacks formed rapidly. However, the larger number of eumelanins resulted in more complex aggregate shapes. This is reflected in the bimodal angle distribution at early times (the black line showing the average over 0-100 ns). The two insets show snapshots at 100 ns (right; T-shaped stacking) and at 2 *μ*s (left). The final equilibrium structure is still a single stack with distribution similar to the smaller systems, but the equilibrium stack formed through formation and subsequent aggregation of smaller stacks that were sometimes perpendicular to each other. Repeated simulations with different initial conditions also showed the same trend as above, see Fig. S1.

### Intermediate to large systems

Figure 3c shows that the system with eight eumelanins starts to show complex aggregation behavior. Another trend is that the equilibrium, or rather steady-state, distributions, albeit maintaining their shape, become broader.

A further increase in system size confirms these observations. Figure 3d shows the angle distribution for a system with 27 eumelanins. The black line (average over 0-100 ns) shows that at early times the distribution is very broad indicating that all orientations are present. When the time approaches 2 *μ*s, the molecules have again stacked and exhibit the characteristic shape and peak close to 15° as the red curve (average over the last 100 ns) displays. The inset shows the system at the end of the simulation at about 2 *μ*s. Instead of a single stack, the system has formed two parallel stacks that are in contact. This shape is stable and was reached also in replica simulations.

Figures 4a–c show the structure and the behavior of water molecules in the vicinity (within 3.5Å) of the two stacks in the system of 27 eumelanins after 1.6 *μ*s. Water molecules are shown as a combination of surface and licorice representations in order to capture the essential features; the water molecules shown using licorice representation are the ones that hydrogen bond (H-bond) with eumelanin molecules. The snapshots show that water is excluded between the eumelanins within the stacks and between the two parallel stacks. Figure 4d shows the distances between the COGs of two eumelanins in purple and the shortest distance between two atoms of neighboring eumelanins in yellow. The interlayer (=shortest) distances were found to be 2.9-3.5 Å which is in good agreement with Meng and Kaxiras^53^ who measured 3.3-3.5 Å in their vacuum DFT calculations. Our results are also in good agreement with Chen et al.^42^ who found, in the absence of water, the interlayer distance to be on average 3.3 Å both in simulations and transmission electron microscopy experiments. We also measured the COG-COG distances and they were typically found to be in the range of 3.5-4.4 Å although larger distances also occur as the angle distributions in Figs. 3a and 3b indicate.

**Figure (4).**
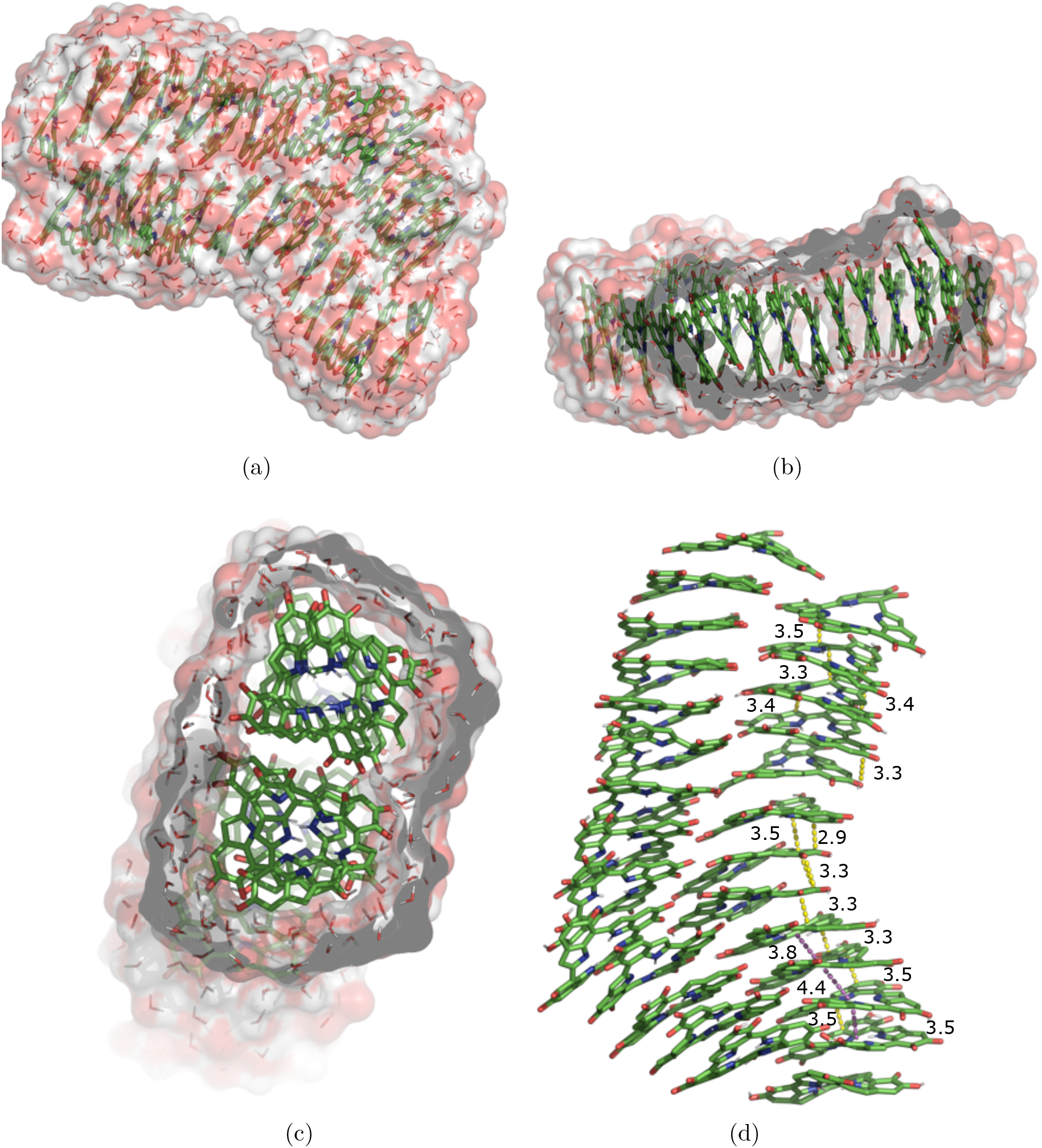
Snapshots of 27 eumelanins in an aqueous solution at 1,600 ns. The eumelanins and also the water molecules with H-bonds to the eumelanins are shown in licorice. The rest of the water molecules are represented using ball and licorice. a) Water molecules within 3.5 Å of eumelanins. b) Sliced image viewed from one of the sides of the system. No water molecules are trapped between the layers. c) Sliced image viewed from the top of the system. The are no water molecules between the two neighboring eumelanin stacks. d) Selected distances between the COGs of two eumelanins are shown in purple and distances between two eumelanins are shown in yellow.

Formation of multiple stacks is a characteristic of large eumelanin systems and to study that, systems with 64 and 343 eumelanins were simulated. The angle distributions behave similarly to the 27 eumelanin system and are not shown. In both cases, several stacks form. Figure 5 shows snapshot from the 343 eumelanin system after 250 ns. As the figure shows, the stacks show complex behavior and they tend to bend and form curved lines. The behavior and scale are very similar to the transmission electron microscopy images of large aggregates in the study of Chen et al.^42^

**Figure (5).**
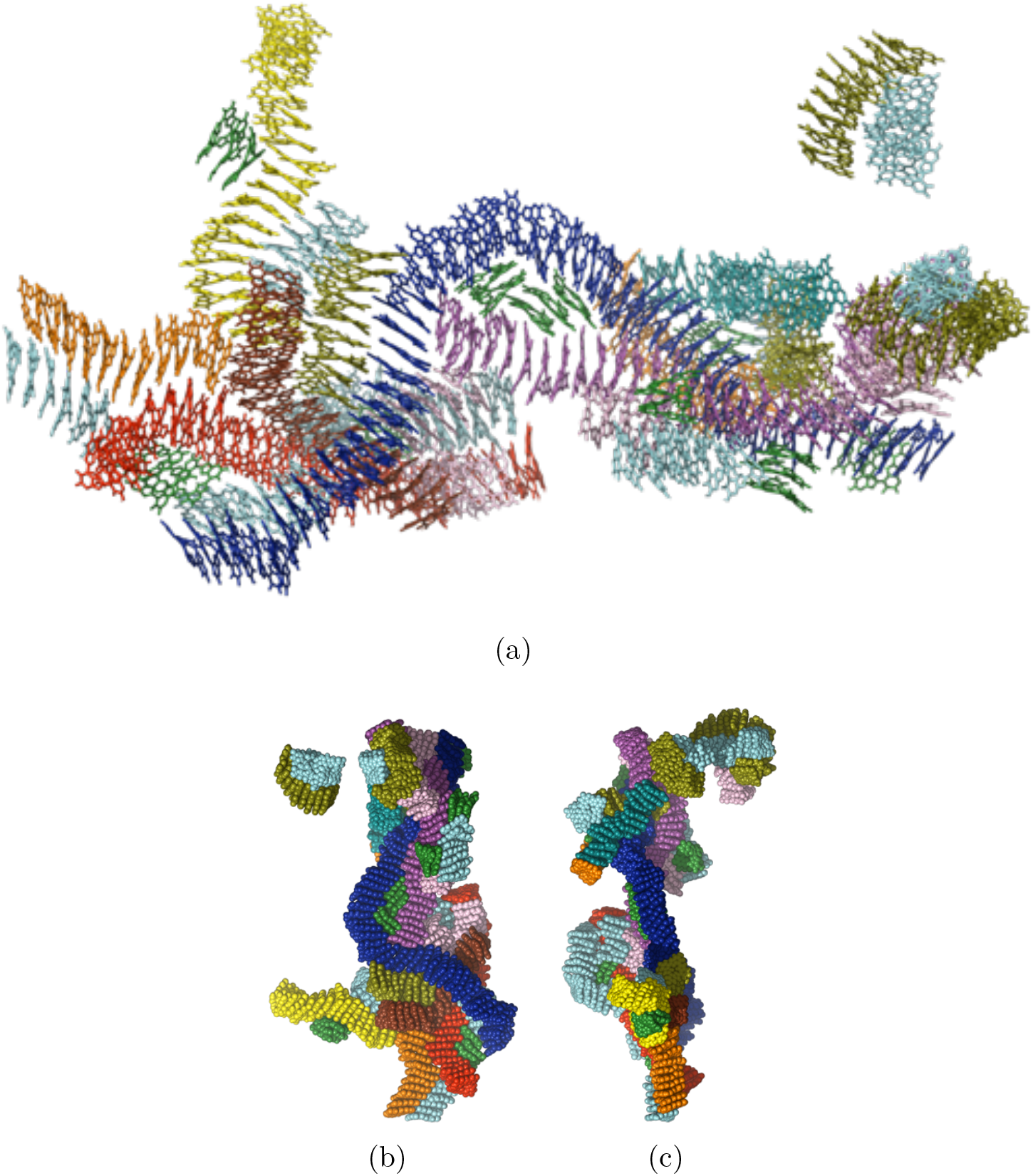
Snapshots of 343 eumelanins in an aqueous solution after 250 ns. Water molecules are not shown for clarity. a) Eumelanins are shown in licorice representation. b) The same using van der Waals representation. c) The same system as in b) but from a different angle. Colors represent different stack sizes: 2 or 3 (green), 4 or 5 (pink), 6 or 7 (cyan), 8 or 9 (olive), 11 (orange), 13 (teal), 20 (yellow), 27 (red), 33 (magenta), and 53 (blue). Comparison with the transmission electron microscopy images in Fig. 3 in Chen et al.^42^ shows very good qualitative agreement.

### Stack size distributions

We analyzed the stack size distributions from single runs. Ideally, distributions would be averaged over a large number of simulations but that was not feasible due to the large system sizes. For systems of two, four and eight eumelanins, the final distribution always consisted of a single stack, this result was also obtained in all replicates (repeated simulations but from different initial conformations), see the black, red and green bars in Figs. 6a and 6b.

**Figure (6).**
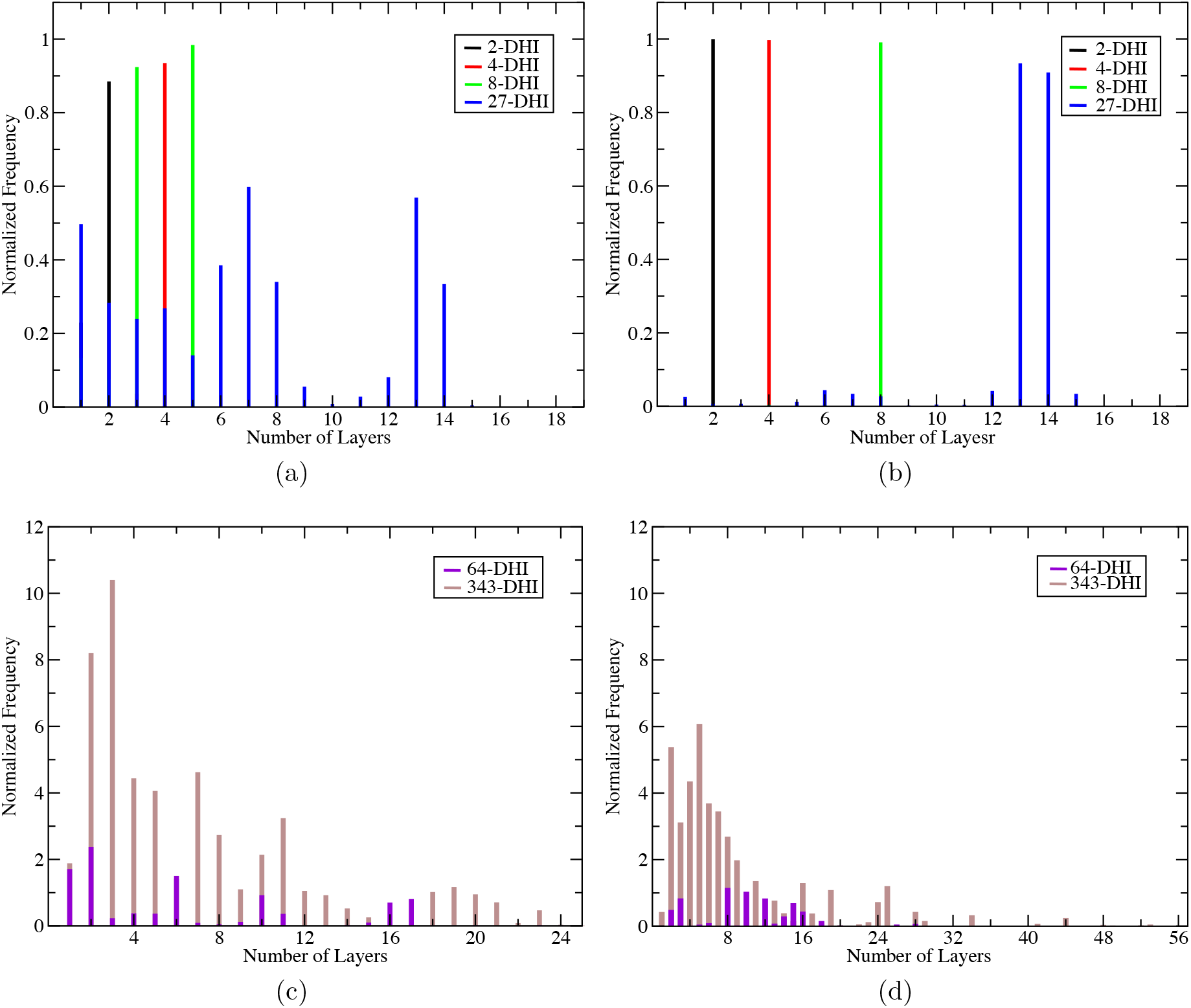
Stack size distributions from single simulation runs. a) Average over the first 100 ns for 2-27 molecules and b) over the last 100 ns. In systems of 2-8 eumelanins, the molecules form a single stack at the end of the simulation. c) Average over the first 100 ns of 64 molecules and over the first 20 ns of 343 molecules. d) Average over the last 100 ns of 64 molecules and over last 20 ns of 343 molecules.

The stack size distributions become more complex when the system size grows. The distribution for the system of size 27 is shown with blue bars in Figs. 6a and 6b. The figures show that at early times the molecules are distributed randomly. At the end (Fig. 6b), however, two approximately equal size stacks are present (see also Fig. 4). The lengths of the two stacks of 13 and 14 molecules are ≈ 46.2 Å and ≈ 48.8 Å, respectively.

For the larger system of 64 eumelanins the most probable stack sizes at the end of simulation are 8 and 10 molecules, and the largest stacks have 15 and 16 molecules Fig. 6c. The lengths of stacks of 3, 8, 10, 12, 15 and 16 molecules correspond approximately to 8.9, 25.0, 35.0, 40.1, 41.6 and 54.9 Å, respectively.

As already observed in Fig. 5, the largest system of 343 eumelanin molecules forms a distribution of nanoscale stacks demonstrating a clear lack of long-range order. The emergence of multiple stacks, bending and disorder are also visible in the smaller system of 27 molecules in Fig. 4. During the first time interval the stacks sizes of the large 64 and 343 systems show the same behavior as the 27 eumelanin simulations, Fig. 6c. At the end, the system has formed stacks of 16, 19, 24, 25 and 53 molecules with corresponding lengths of 53.5, 64.3, 78.9, 73.8 and 139.9 Å as shown in Fig. 6d.

### Radial Distribution Functions (RDF)

The water-eumelanin RDFs in Fig. 7a show that as eumelanin concentration increases, water becomes pushed further away from the eumelanin molecules. This is consistent with aggregation behavior. All of the RDFs in Fig. 7a have a small rise at around 8 Å. This due to water hydrogen bonding to eumelanin’s carbonyl oxygens in the IQ and MQ groups (see Fig. 1b).

**Figure (7).**
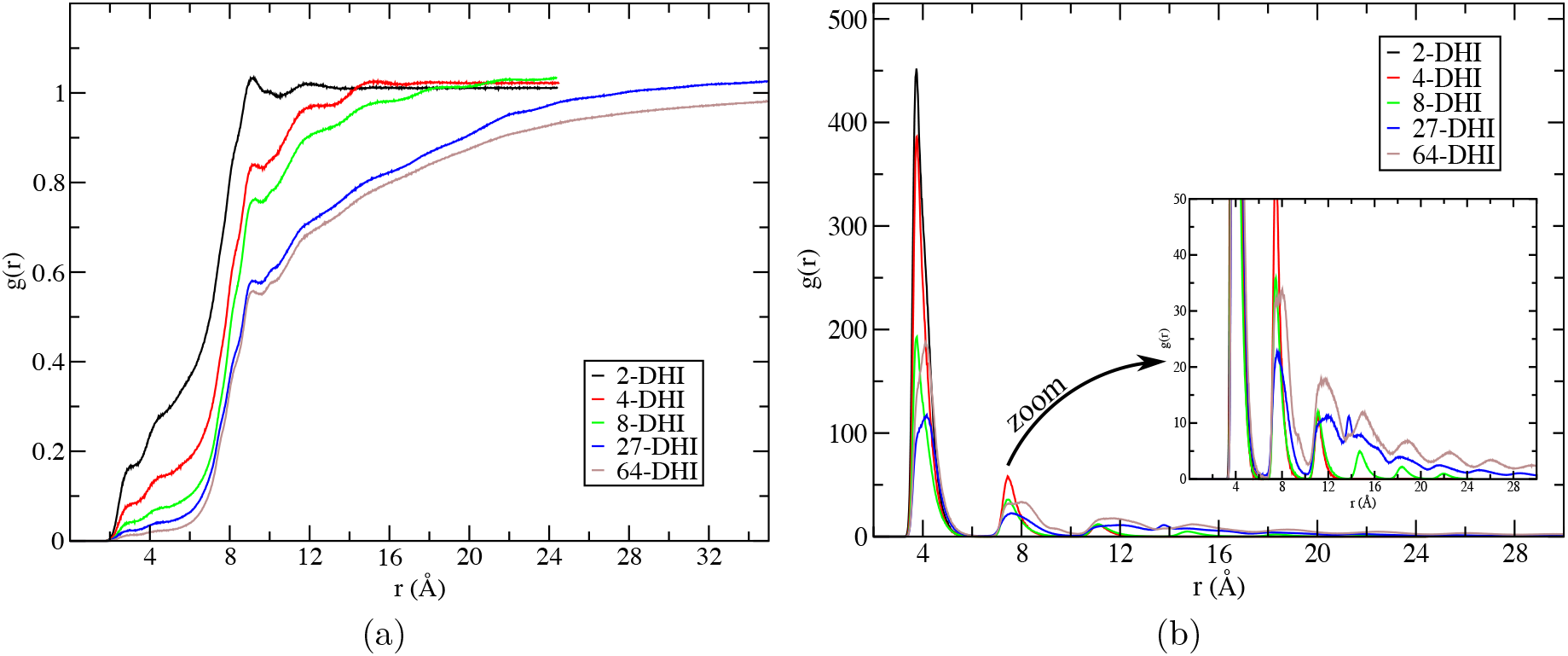
Radial distribution functions. a) RDF between water molecules and eumelanins and b) RDF between eumelanin centers.

The eumelanin-eumelanin RDFs in Fig. 7b show their tendency to stack at the intermolecular distance of 3.75 Å in systems of 2, 4 and 8 eumelanins, and at 4.15 Å in the cases of 27 and 64 eumelanins. The two eumelanin system has only a single peak at 3.75 Å, but the rest of systems show second peaks at around 7.45 Å and higher order peaks at longer distances. These peaks and their broadening result from short-range ordering along the stacks as well as from the emergence of parallel stacks and T-shaped configurations (see Fig. 5).

### Hydrogen Bonding

Figures 8 and 9 show the time evolution of the number of water-eumelanin H-bonds and the running average for the systems of 2-27 (Fig. 8), and 64 and 343 (Fig. 9) eumelanins. To define an H-bond, an angle < 30° and a donor-acceptor distance < 3.5 Å were used as the criteria.

**Figure (8).**
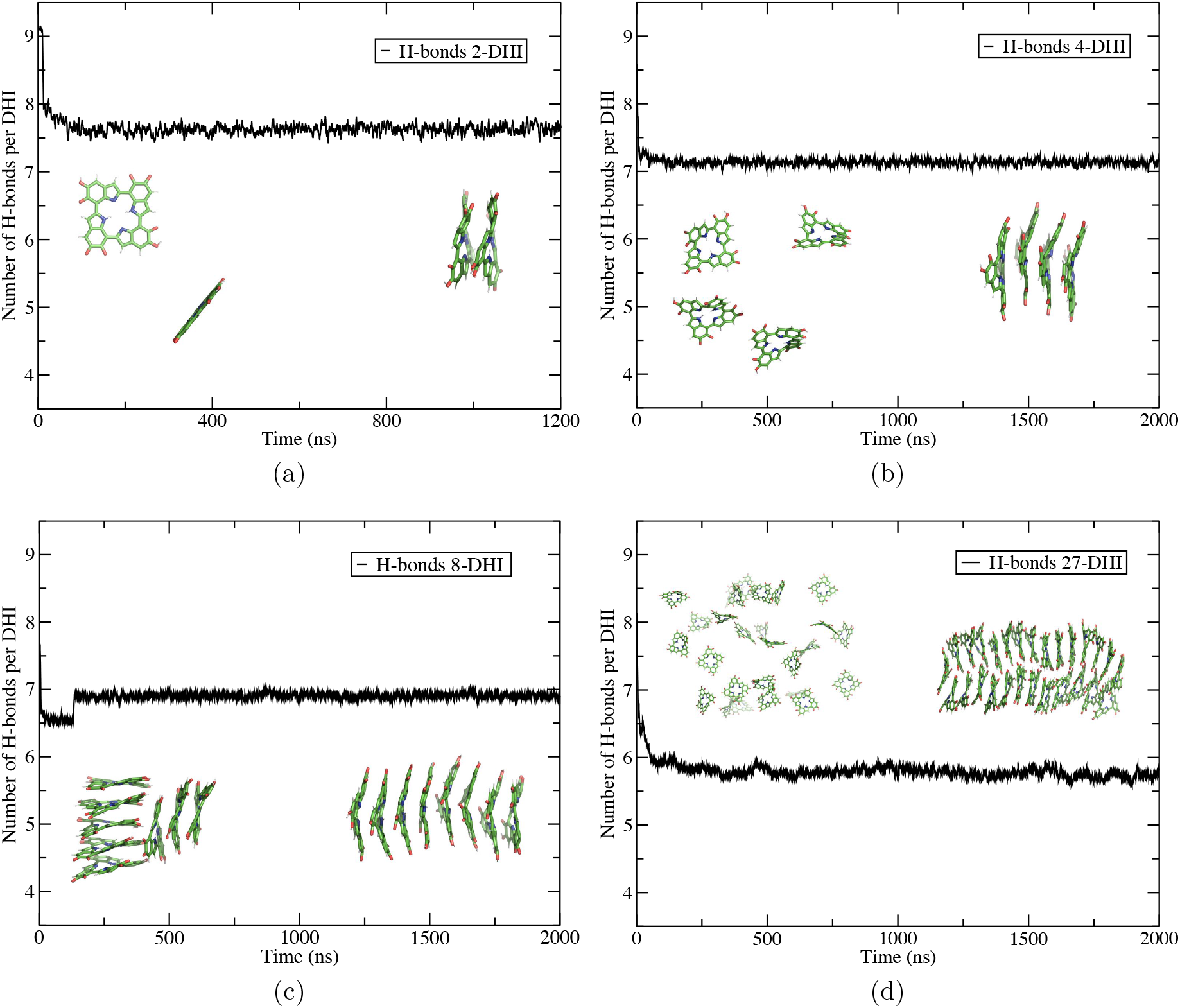
Time evolution (running average over 250 ps) of the number of hydrogen bonds with water per eumelanin. a) System of two eumelanins. The inset shows the random initial configuration and final state after 1.2 *μs*. b) Four eumelanins. Snapshots show the initial and final configurations. c) Eight eumelanins. d) 27 eumelanins. As a general trend, the number of H-bonds/eumelaning decreases when the system size increases. This is due to the fact (see Fig. 4) that water is expelled from the layers and between the stacks.

**Figure (9).**
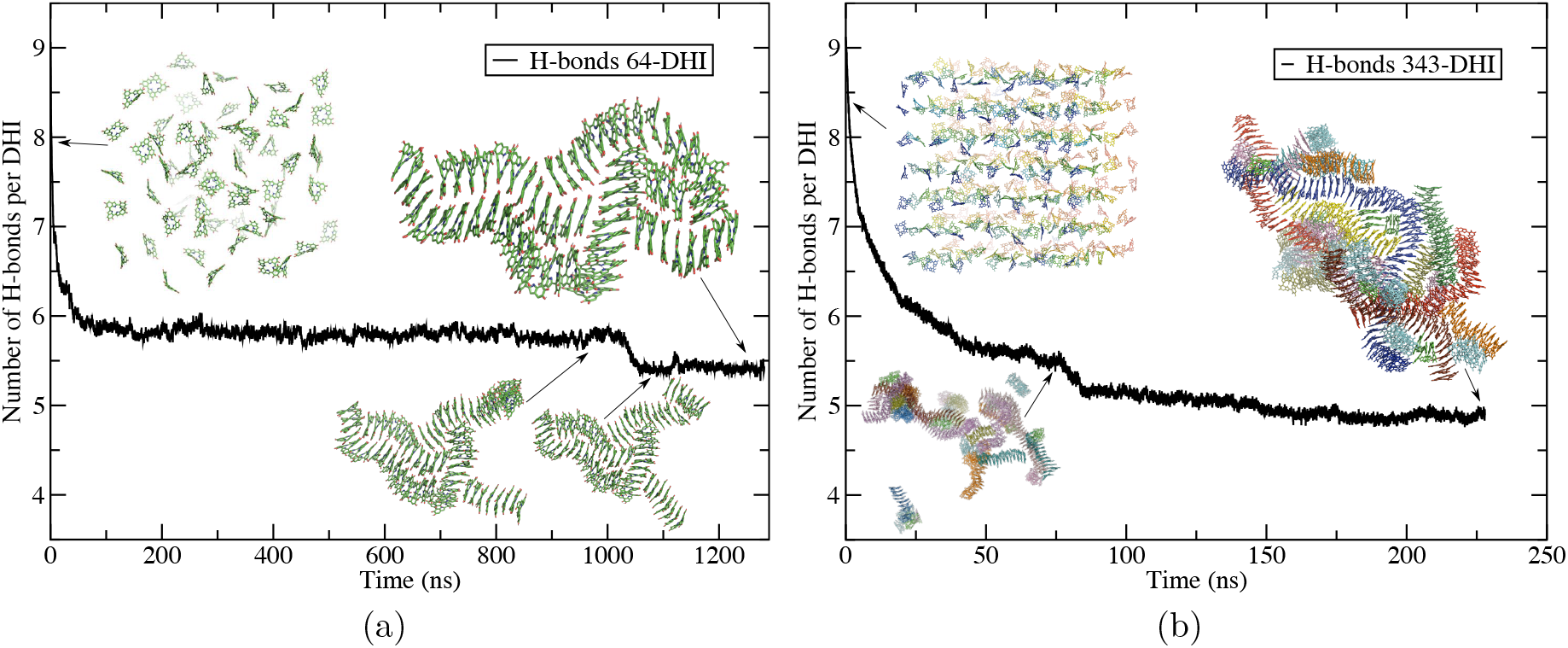
Time evolution of the number of eumelanin-water hydrogen bonds per eumelanin in systems of 64 and 343 eumelanins. The inset shows snapshots at different times during the simulations.

As Fig. 8 shows, the average number of H-bonds per eumelanin decreases from 7.5 to 4.9 when the number of eumelanins increases from two to 343. This is due to the fact that each eumelanin has four building blocks and the number of H-bonds per building block depends on the size of the aggregate and ranges from 1.9 to 1.2. The result is in good agreement with Supakar et al.^46^ who showed that the number of H-bond decreases when the number of melanin building blocks increases. That means that eumelanins favor stacking interactions and, interestingly, that they favor to stay in one layer and plane as observed in insets of Figs. 8 and 9a.

### Free Energy of Stacking

To evaluate the stacking free energy, the potential of mean force (PMF) as a function of the distance between the COMs of two eumelanins was calculated. Four different systems with two parallel eumelanins were studied such that a) the molecules had the exact same orientation, b) the same orientation but one molecule rotated by 90°, c) mirrored or flipped with respect to each other and d) mirrored and rotated by 90°. The systems are shown in Fig. 2 and the corresponding PMFs in Fig. 10. The minimum for all the systems except the one with 90° rotation occurs at 3.6 Å. The one with 90° rotation has its free energy minimum at the distance of 3.7 Å as shown in Fig. 10.

**Figure (10).**
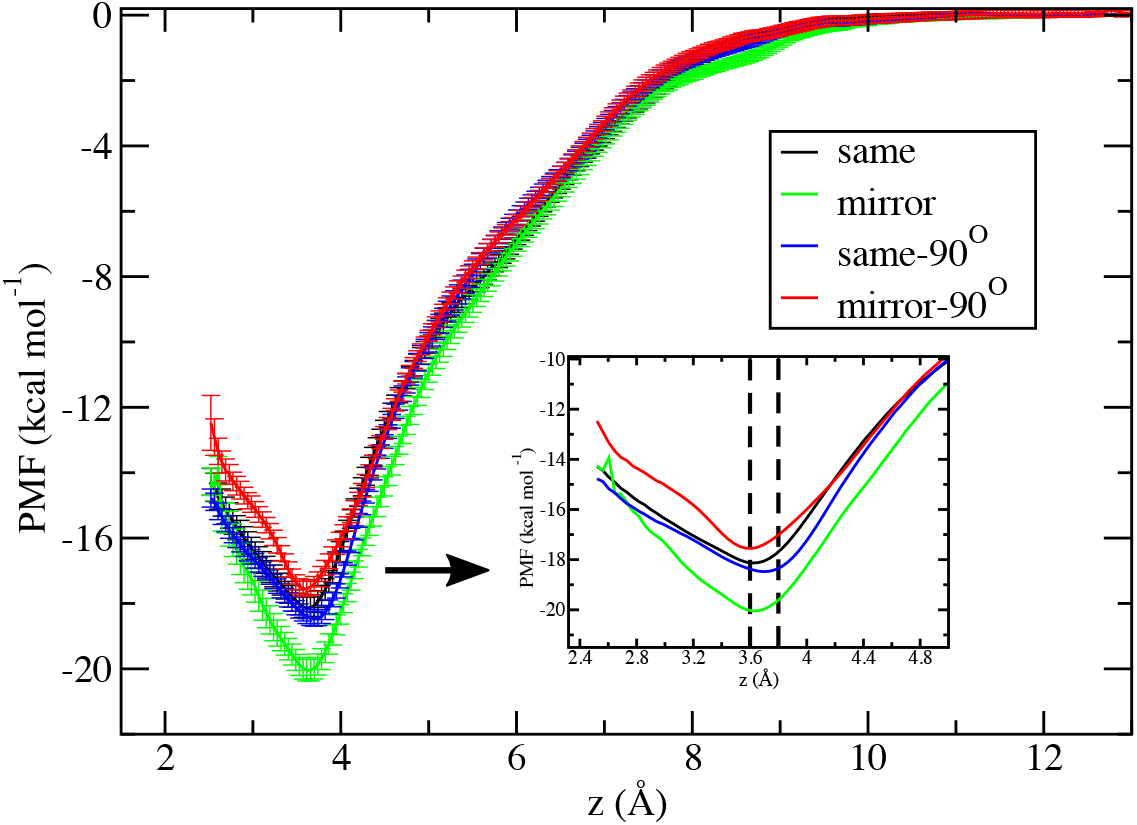
PMFs of two parallel eumelanins. See Fig. 2 for the definitions for rotation and mirroring. The inset shows a close-up of the range where the minima are located.

### Eumelanin Dimers Evaluated by DFT

To better understand stacking, quantum mechanical DFT calculations were carried out on various dimeric orientations. Six dimeric conformations were optimized, and their energies were compared. The Stacked-1 was made by placing one eumelanin on top of the other so that the former was eclipsed. This conformation corresponds to the structure shown in Fig. 2a. There is a 90° rotation in the case of Stacked-2. In Planar-1 conformer, the MQ units of the neighboring eumelanins are facing each other and Planar-2 is made by putting the MQ unit of one eumelanin facing the IQ unit of the neighboring eumelanin. There are about 40° and 80° differences between the eumelanin planes in the Twisted and T-shape conformations, respectively. There are also slight bends in the eumelanin plane regardless of the dimeric conformation. A few other possible dimeric conformations did not meet the optimization criteria and hence were discarded.

As predicted by all of our DFT calculations (Table 2) the Stacked-2 conformation is the most stable one among all the considered conformers. In fact, both Stacked-1 and Stacked-2 have considerably lower energies in comparison with all the other conformations. It can be concluded that stacking energy (*π* – *π* interaction) of the two eumelanins is much larger than the electrostatic interactions (hydrogen bonds) in Planar-1, Planar-2, and Twisted conformers. The T-shaped conformation is more stable than the twisted and planar ones, and less stable compared to the stacked conformations. This is in line with the angle distribution of eumelanin obtained with MD simulations, where the angles corresponding to the stacked conformations (0 – 20°) and to the T-shaped ones (≈ 90°) are the most populated ones (see Fig. 3).

**Table (2).** Relative stabilities of eumelanin dimers (kcal/mol) calculated using a polarizable continuum model (IEF-PCM) at different levels of theory. Optimizations were carried out at IEF-PCM-CAM-B3LYP/6-311G(d,p) level. See Fig. 11 for the orientations.

|  | M06-2X/<br>6-311++G(2df,2pd) | CAM-B3LYP-D3BJ/<br>6-311++G(2df,2pd) | B3LYP-D3BJ/<br>6-311++G(2df,2pd) |
| --- | --- | --- | --- |
| Twisted | 0.0 | 0.0 | 0.0 |
| Planar-1 | -0.3 | -1.3 | -1.4 |
| Planar-2 | -2.1 | -3.4 | -3.7 |
| T-shape | -4.5 | -6.9 | -7.5 |
| Stacked-1 | -16.0 | -25.2 | -26.3 |
| Stacked-2 | -21.9 | -29.3 | -31.3 |

Our results are in line with the findings of Barone et al.^84^ on simple chatechol dimers, where sheared stacked conformations were found to be the most stable arrangement obtained at DFT level. In addition, MP2 calculations by Prampolini and coworkers^85^ have shown that pyrole rings such as the IQ and MQ subunits of eumelanin form similar sheared stacked dimers, where both stacking and hydrogen bond (electrostatic) interactions contribute to the stability of the dimer. Even in smaller systems such as pyroles edge-to-edge dimers formed by hydrogen bonds are energetically less favourable and stacking interaction contribution provides larger stability in stacked forms. Stability of the T-shaped structures lie between the stacked and edge-to-edge dimers.

We would like to note that only a handful of dimeric conformations were examined here, and the effects of water were only considered through implicit solvent models. Hence, a quantitative comparison between the QM (this section) and MD results is not practical. The potential of mean force for a parallel staggered geometry eumelanin-eumelanin pair is in the range of −17 to −20 kcal/mol which is in good agreement with M06-2X/6-311++G(2df,2pd).

To elucidate the behaviour of the dimers, the electrostatic potential map (ESP Map) of the eumelanin molecule was calculated and is depicted in Fig. 12. The isodensity surface of 0.02 a.u is appropriate for the analysis of non-covalent interactions as it roughly corresponds to the surface that two non-reacting molecules can approach. The electrostatic potential map shows three areas: First, the porphyrin-like core shows as a red surface indicating electron-rich nature. Second, the green color indicates the presence of positive charges, and finally the periphery is in yellow (electron rich). This means that when two eumelanin molecules are stacked on top of one another, the repulsion due to the positive core may outweigh any attractive forces unless there is a rotation and possibly a shift: The parallel staggered geometry (Stacked-2) is preferred over face-to-face (Stacked-1), planar or edge-to-face (T-shaped) geometries, see also Table 2.

A different scenario based on the eumelanin structure has been previously proposed by Meng et al.^53^ They suggested that covalent bonds may be formed between two stacked eumelanins and that the resulting structures can explain the optical properties of eumelanin aggregates. Although according to their calculations fused structures are more stable than two individual eumelanin molecules, they did not provide energy barriers for formation of such structures. Such covalent bonds between two eumelanins would disturb the delocalized electrons over the molecular plane, perturb the aromaticity of the molecule, and hence alter its stacking properties. In addition, formation of a fused structure would require one molecule to be completely on top of the second one (eclipsed), which according to our EPM calculation (Fig. 12) would not be energetically favourable. Such eclipsed structures were not observed in our DFT calculations (Fig. 11). Although comparison between stacking properties, the behaviour of fused tetramers (as proposed by Meng et al.^53^) and the behaviour of single eumelanins would potentially provide useful information, such a study is beyond the scope of the current work.

**Figure (11).**
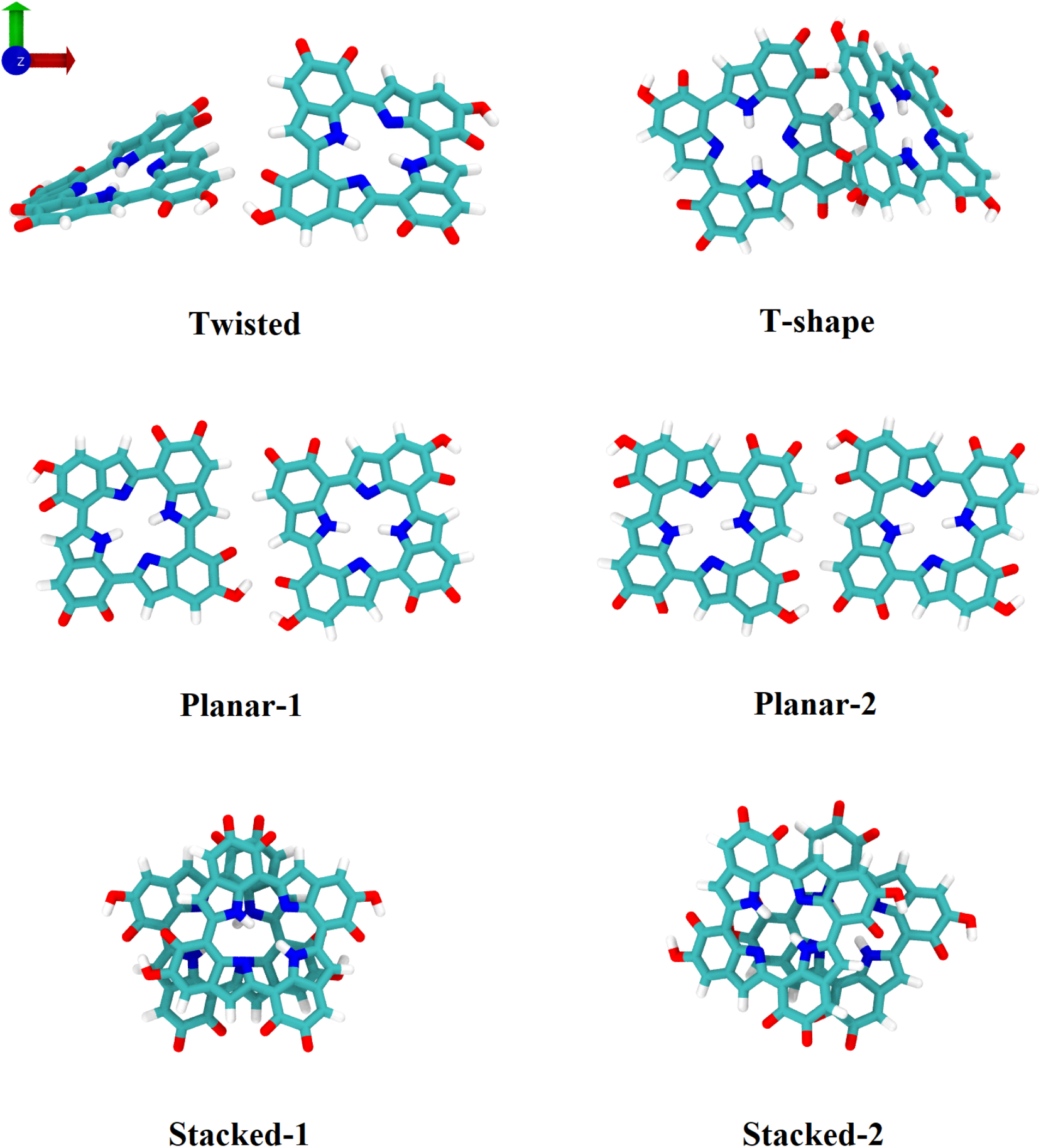
Different eumelanin dimer conformations considered with DFT. There is an angle of 40° and 80° between the eumelanin monomers in the Twisted and T-shape conformations, respectively. In the Planar conformations the dimers the two eumelanins share a plane, where in Planar-1, two MQ and two IQ subunits from the neighboring eumelanins are facing each other, whereas in Planar-2, one MQ of eumelanin is facing an IQ subunit of the second eumelanin. The monomers in the Stacked conformations are nearly parallel. In Stacked-1 a eumelanin is flipped (by 90°) and in Stacked-2 the two eumelanins are mirror images.

**Figure (12).**
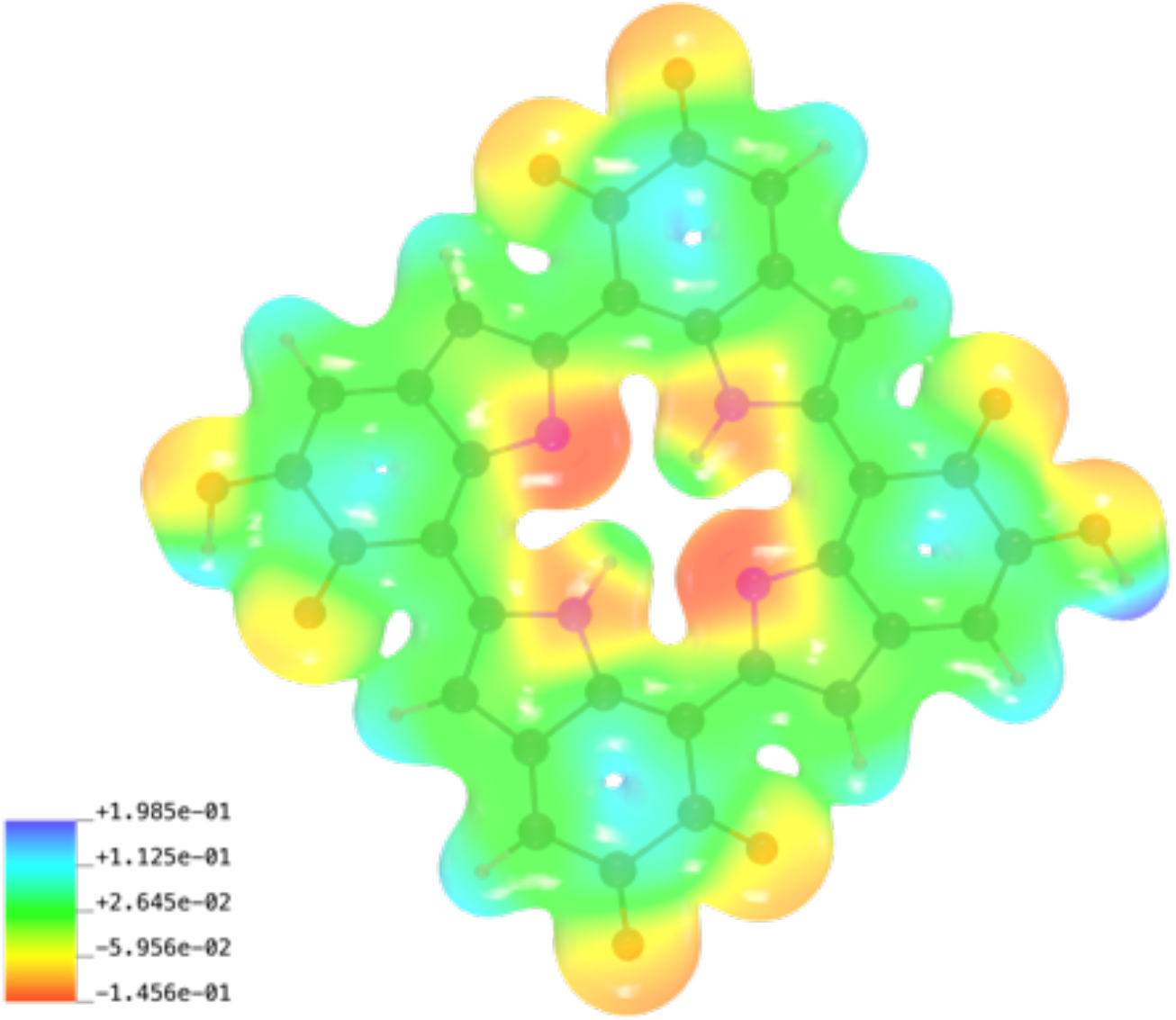
Electrostatic potential map of an eumelanin molecule. Electrostatic potentials are mapped on the surface of the electron density of the 0.02 unit. The red surface corresponds to a region of negative electrostatic potential, whereas the blue color corresponds to the positive potential. The numbers of positive and negative potentials are given in atomic units of length cubed 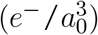.

## Conclusions

In this study, we have examined eumelanin aggregation using three different methods: 1) multi-*μs* long classical MD simulations, 2) umbrella sampling for free energy calculations and 3) DFT calculations for the detailed physical mechanisms of eumelanin stacking, as well as parametrisation of the DHI eumelanin (Table S1).

The classical MD simulations were performed in aqueous solutions at different concentrations in systems consisting of 2, 4, 8, 27, 64, and 343 DHI eumelanin molecules. All systems showed aggregation and stacking with interlayer distance between the eumelanin molecules being less than 3.5 Å. Our results show that upon aggregation, the average number of hydrogen bonds between the individual eumelanins and solvent decreases. This change is more profound at higher concentrations of eumelanins and directly related to changes in the domain size and aggregate structure (see Figs. 8, 9 and S1). The distribution of angles along the stacks shows a semi-bimodal distribution (see Fig. 3). This indicates the preference of eumelanins to form stacked and perpendicular (T-shaped) bundles. The abundance of stacked and T-shaped conformations observed in MD simulations is determined by their lower energies (i.e., more stable) relative to other conformations such as planar or twisted as confirmed by our DFT calculations (see Fig. 11 and Table 2). Stacking has been previously reported for eumelanin in the absence of water or in nearly dry system,^42,47^ and also observed in transmission electron microscopy experiments^42^.

In both MD simulations and DFT calculations, stacked eumelanins tend to form sheared stacks and, consequently, the molecules do not completely eclipse one another (see Fig. 11, Stacked-1 and 2). This deviation from fully eclipsed conformation may play a role in formation of curving of the nanoscale domains observed for the large stacks (Figs. 4 and 5). The appearance of sheared conformations between the two eumelanin is well-explained by the electrostatic potential map of the molecule (Fig. 12). The negative potential at the center of the molecular plane and its surrounding positive potential disfavours fully eclipsed conformations and on the other hand, a sheared stack allows two DHIs to form favorable attractive interactions.

## Supporting Information

Table S1: Atom types, coordinates (in nm) and ESP charges of DHI eumelanin optimized using DFT calculations. Figure S1: Time evolution (running average over 250 ps) of the number of hydrogen bonds with water per eumelanin for a system of eight eumelanins. The inset shows snapshots of the system at 100 ns, 1.1 *μ*s, 1.7 *μ*s, 2.1 *μ*s and the final state after 2.7 *μ*s.

## Acknowledgement

MK acknowledges the financial support by the Natural Sciences and Engineering Research Council of Canada (NSERC) and Canada Research Chairs Program. CGTF is a recipient of the Western University Postdoctoral Fellowship Award. Computational resources were provided by Compute Canada and SharcNet.

## Graphical TOC Entry

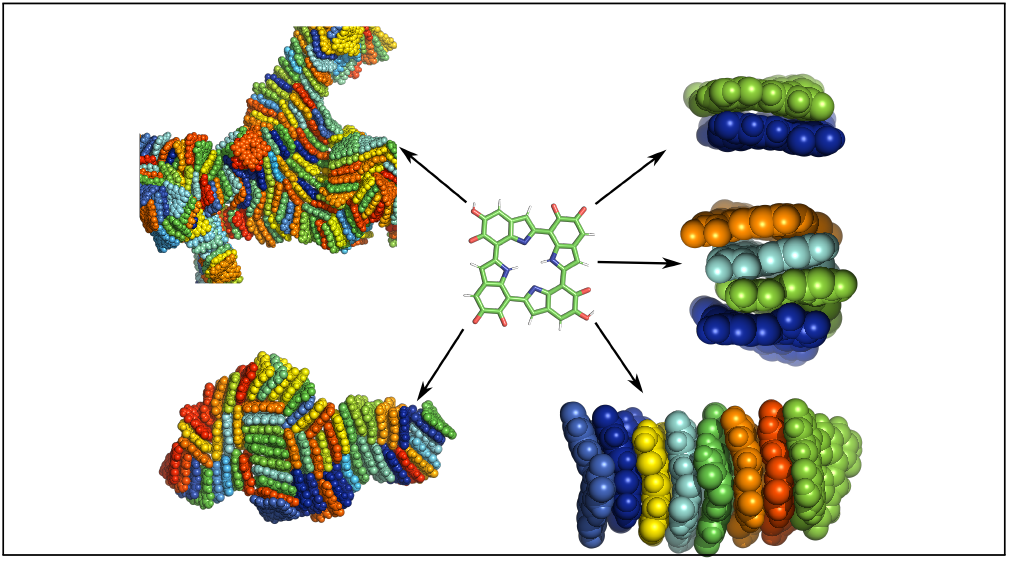

