## Supplementary Information for "Free energy and stacking of eumelanin nanoaggregates"

Table S1: Atom types, coordinates (in nm) and ESP charges of DHI eumelanin optimized using DFT calculations.

| | Atom Name | X(nm) | Y (nm) | Z(nm) | ESP charge |
|---|---|---|---|---|---|
| 1 | H | 1.038 | 2.106 | 1.000 | 0.4565 |
| 2 | O | 1.032 | 2.008 | 1.000 | -0.5393 |
| 3 | C | 1.155 | 1.955 | 1.000 | 0.1850 |
| 4 | C | 1.272 | 2.024 | 1.000 | -0.3765 |
| 5 | C | 1.153 | 1.804 | 1.000 | 0.6395 |
| 6 | H | 1.274 | 2.133 | 1.000 | 0.1998 |

Continued on next page

Table S1 – continued from previous page

| | Atom Name | X(nm) | Y(nm) | Z(nm) | ESP charge |
|---|---|---|---|---|---|
| 7 | C | 1.395 | 1.950 | 1.000 | 0.0039 |
| 8 | O | 1.044 | 1.750 | 1.000 | -0.4366 |
| 9 | C | 1.283 | 1.728 | 1.000 | -0.4581 |
| 10 | C | 1.401 | 1.802 | 1.000 | 0.3803 |
| 11 | C | 1.524 | 1.992 | 1.000 | -0.3127 |
| 12 | C | 1.275 | 1.584 | 1.000 | 0.3062 |
| 13 | N | 1.532 | 1.759 | 1.000 | -0.4970 |
| 14 | H | 1.561 | 2.094 | 1.000 | 0.2382 |
| 15 | C | 1.607 | 1.871 | 1.000 | 0.4547 |
| 16 | C | 1.164 | 1.503 | 1.000 | -0.4309 |
| 17 | N | 1.390 | 1.501 | 1.000 | -0.0519 |
| 18 | C | 1.750 | 1.875 | 1.000 | -0.3805 |
| 19 | H | 1.061 | 1.537 | 1.000 | 0.2509 |
| 20 | C | 1.208 | 1.368 | 1.000 | 0.1862 |
| 21 | H | 1.486 | 1.537 | 1.000 | 0.1676 |
| 22 | C | 1.356 | 1.371 | 1.000 | 0.1101 |
| 23 | C | 1.824 | 2.005 | 1.000 | 0.4885 |
| 24 | C | 1.831 | 1.760 | 1.000 | 0.1157 |
| 25 | C | 1.136 | 1.252 | 1.000 | -0.3597 |
| 26 | C | 1.437 | 1.256 | 1.000 | -0.3973 |
| 27 | O | 1.771 | 2.114 | 1.000 | -0.4552 |
| 28 | C | 1.981 | 2.007 | 1.000 | 0.4706 |
| 29 | N | 1.797 | 1.630 | 1.000 | -0.0675 |
| 30 | C | 1.980 | 1.763 | 1.000 | 0.1537 |
| 31 | H | 1.027 | 1.250 | 1.000 | 0.2060 |
| 32 | C | 1.206 | 1.124 | 1.000 | 0.4728 |
| 33 | C | 1.580 | 1.259 | 1.000 | 0.4855 |
| 34 | C | 1.363 | 1.125 | 1.000 | 0.4848 |
| 35 | O | 2.037 | 2.114 | 1.000 | -0.4196 |
| 36 | C | 2.051 | 1.879 | 1.000 | -0.3563 |
| 37 | H | 1.701 | 1.594 | 1.000 | 0.1677 |
| 38 | C | 1.911 | 1.547 | 1.000 | 0.3030 |
| 39 | C | 2.023 | 1.627 | 1.000 | -0.4031 |
| 40 | O | 1.149 | 1.017 | 1.000 | -0.4170 |
| 41 | N | 1.655 | 1.372 | 1.000 | -0.5309 |
| 42 | C | 1.663 | 1.138 | 1.000 | -0.2790 |
| 43 | O | 1.416 | 1.016 | 1.000 | -0.4452 |
| 44 | H | 2.160 | 1.881 | 1.000 | 0.2027 |
| 45 | C | 1.903 | 1.403 | 1.000 | -0.4507 |
| 46 | H | 2.126 | 1.593 | 1.000 | 0.2451 |
| 47 | C | 1.786 | 1.328 | 1.000 | 0.4442 |
| 48 | H | 1.626 | 1.036 | 1.000 | 0.2232 |

Continued on next page

**Table S1 – continued from previous page**

| | Atom Name | X(nm) | Y(nm) | Z(nm) | ESP charge |
|---|---|---|---|---|---|
| 49 | C | 1.792 | 1.180 | 1.000 | -0.0971 |
| 50 | C | 2.031 | 1.326 | 1.000 | 0.5794 |
| 51 | C | 1.915 | 1.106 | 1.000 | -0.2841 |
| 52 | O | 2.143 | 1.378 | 1.000 | -0.4878 |
| 53 | C | 2.031 | 1.176 | 1.000 | 0.1747 |
| 54 | H | 1.915 | 0.997 | 1.000 | 0.2227 |
| 55 | O | 2.152 | 1.122 | 1.000 | -0.4981 |
| 56 | H | 2.216 | 1.197 | 1.000 | 0.4129 |

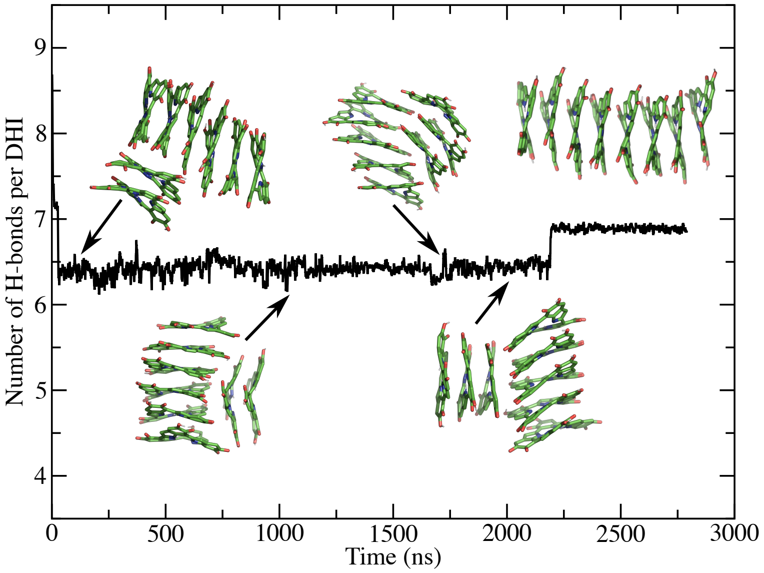


Figure S1: Time evolution (running average over 250 ps) of the number of hydrogen bonds with water per eumelanin for a system of eight eumelanins. The inset shows snapshots of the system at 100 ns, 1.1 $\mu$s, 1.7 $\mu$s, 2.1 $\mu$s and the final state after 2.7 $\mu$s.
